# Evolutionary history and polyploidization lead to rapid shifts in chemodiversity of *Hypericum*

**DOI:** 10.64898/2026.08.18.744459

**Authors:** Xue Xiao, Rabea Schweiger, Eric Ron Stein, Thomas Dussarrat, Marcus A. Koch, Caroline Müller

## Abstract

- Polyploidization can profoundly affect plant metabolite biosynthesis, yet its influence on chemodiversity remains poorly understood, despite the central role of chemodiversity in mediating plant interactions with the environment. The coexistence of facultative apomictic and sexual reproductive systems across ploidy levels in *Hypericum* provides an excellent model for investigating the evolution of chemodiversity following polyploidization.
- We analyzed ploidy levels and leaf metabolic fingerprints across selected populations of three *Hypericum* taxa, *H. maculatum*, *H. perforatum* subsp. *perforatum* and *H. perforatum* subsp. *veronense*.
- Polyploidization was common across all three taxa. Leaf metabolic fingerprints were more pronouncedly differentiated by the ploidy level of the mother plant (F0) than that of the offspring (F1). Although unique metabolic features emerged in plants of most ploidy levels, diploid plants exhibited fewer metabolic features than polyploid plants. Higher Shannon diversity, functional Hill diversity, and intensities of features belonging to specific chemical families were associated with higher F0 ploidy levels in *H. perforatum* subsp. *perforatum*, but not in *H. maculatum* and *H. perforatum* subsp. *veronense*.
- Our findings demonstrate that polyploidization can lead to rapid shifts in chemodiversity across generations in *Hypericum*. The fast divergence in chemodiversity associated with polyploidization in *H. perforatum* may contribute to its remarkable invasive potential.

## Introduction

Polyploidization, defined as an increase in whole-genome copy number, is widespread in plants and is estimated to occur in more than 35% of angiosperm genera (Wood *et al*., 2009). Polyploidization can have profound consequences for plant metabolism by altering gene dosage, regulatory networks, and biosynthetic pathways (Gaynor *et al*., 2020). Accordingly, polyploidization has been shown to enhance the production of key specialized metabolites, for example, artemisinin in root cultures of *Artemisia annua* (Asteraceae) and terpenoids in *Thymus persicus* (Lamiaceae) (Jesus-Gonzalez & Weathers, 2003; Tavan *et al*., 2015). While these studies demonstrate effects of polyploidization on the production of individual metabolites or chemical families, its impact on overall plant chemodiversity remains poorly understood. Moreover, the mode of polyploidization may shape plant chemodiversity in different ways. In particular, allopolyploidization, compared to autopolyploidization, may foster metabolite diversification, because the fusion of divergent genomes may enable novel metabolic pathways, enzyme combinations, and increased heterozygosity, which in turn may generate new chemical profiles (Soltis *et al*., 2014; Madani *et al*., 2021).

Chemodiversity is vital for plant evolution, as it affects ecological interactions across trophic levels and environments, for example, by contributing to niche differentiation in plants (Müller & Junker, 2022). Increasing evidence highlights that chemodiversity of specialized metabolites shapes multitrophic interactions and microbial assemblies in plant-associated communities (Wetzel & Whitehead, 2020; Petrén *et al*., 2024; Malacrinò *et al*., 2025). For example, higher heterogeneity in the chemodiversity of *Tanacetum vulgare* (Asteraceae) increases pollinator abundance while reducing herbivore pressure (Sasidharan *et al*., 2024; Ojeda-Prieto *et al*., 2025). Similarly, elevated floral scent chemodiversity is associated with higher flower-visitor richness but lower bacterial richness on flowers of different plant communities (Hanusch *et al*., 2025). Beyond ecological functions, different plant metabolites have long been central to human healthcare through the use of medicinal plants for thousands of years. Higher diversity in specialized metabolites increases the likelihood of discovering bioactive compounds with pharmaceutical potential (Madani *et al*., 2021). Understanding the mechanisms that generate and maintain chemodiversity is therefore essential not only for elucidating plant responses to environmental challenges, but also for improving the discovery of medically relevant natural products.

To date, several ecological and evolutionary hypotheses have been proposed to explain the remarkable chemodiversity of plants, including the coevolutionary arm race hypothesis, which posits that plant chemodiversity accumulates in a stepwise evolutionary process driven by the coevolution between plants and their antagonists (Speed *et al*., 2015). By contrast, the interaction diversity hypothesis postulates that chemodiversity arises from distinct selection pressures (Wetzel & Whitehead, 2020). However, both hypotheses primarily emphasize the role of long-term evolutionary drivers and ecological processes in shaping plant chemodiversity (Wood *et al*., 2009; Wetzel & Whitehead, 2020). Beyond that, chemodiversity may also evolve over relatively short time scales through genomic changes, for instance polyploidization (Lichman *et al*., 2020). Accurate measurement of chemodiversity is a prerequisite for understanding both evolutionary origins and ecological functions of chemodiversity. Functional Hill diversity provides a comprehensive characterization of plant chemodiversity, as it integrates metabolite richness, evenness, and structural dissimilarity into a unified framework (Petrén *et al*., 2023). This measure showed positive relationships between chemodiversity and phenotypic plasticity in response to drought and herbivory (Xiao *et al*., 2025), highlighting a possible link between metabolic complexity and responses to environmental challenges.

*Hypericum perforatum* (Hypericaceae) is a perennial plant species native to Western Europe, Asia, and North Africa, which has become widely naturalized across temperate regions worldwide (Molins *et al*., 2014). It has been a well-known medicinal plant for centuries and produces diverse metabolites, such as phenolic acids, flavonoids, terpenoids, and phloroglucinols, which can have antidepressant, anti-inflammatory, antimicrobial, or antiviral activities (Barnes *et al*., 2001, Ghasemi Pirbalouti *et al.,* 2014; Silva *et al.,* 2005). Among *H. perforatum* and the closely related *H. maculatum* (Scheriau *et al*., 2017), three major taxa have been identified based on nuclear markers and plastid genome variation, corresponding to the informal *H. perforatum* subsp. *perforatum, H. perforatum* subsp. *veronense*, and *H. maculatum* lineages (Koch *et al*., 2013). All three taxa contain predominantly sexual diploid populations as well as facultative pseudogamous apomictic polyploid lineages. Polyploid populations commonly include triploid, tetraploid, and hexaploid cytotypes (Koch *et al*., 2013). In Central Europe, natural populations show substantial secondary gene flow between the *H. perforatum* and *H. maculatum* gene pools, and this genetic set-up is also found in regions such as North America where *H. perforatum* is invasive (Molins *et al*., 2014). Previous studies have reported contrasting effects of polyploidization on metabolite composition in *H. perforatum*. For instance, hypericin content has been observed to be higher in diploid than in tetraploid individuals, whereas chloride-induced polyploidization has been associated with increased total phenolic content (Koperdáková *et al*., 2007; Manteghi Tafreshi & Mohammadhassan, 2025). These findings suggest that polyploidization may differentially influence specific metabolite classes. The prevalence of polyploidization, together with a mixed reproductive system, makes the *Hypericum* complex an ideal model for investigating whether polyploidization modulates plant chemodiversity intra- and interspecifically.

To disentangle the effects of ploidy level on leaf metabolic fingerprints, chemodiversity, feature richness, and the intensities of key chemical families, we analyzed leaves of *H. maculatum* and *H. perforatum* (subsp. *perforatum*, and subsp. *veronense*) grown under common climate chamber conditions. We determined both mother plant (F0) and offspring (F1) ploidy levels and assessed their associations with the overall leaf metabolic fingerprint and chemodiversity metrics, i.e., Shannon diversity, functional Hill diversity, and richness, as well as total intensities of phenolic acids, flavonoids, terpenoids, and phloroglucinols, which turned out to be most prominent in our analysis. We hypothesized that leaf metabolic fingerprints are more pronouncedly discriminated by the ploidy level of the offspring than that of the F0 mother plants. In addition, we expected polyploidization to promote the formation of “unique” metabolic features, leading to increased diversity and richness across all three taxa. Finally, we predicted that polyploidization would enhance the biosynthesis of phenolic acids, flavonoids, terpenoids, and phloroglucinols.

## Materials and Methods

### Study species and sampling design

Seeds of *Hypericum* spp. (Hypericaceae) were collected in 2013 from 10 geographically distinct populations, following Koch et al. (2013) (Table 1, Fig. S1). Each population was represented by three F0 mother plants, except for one population (pop320), which was represented by only two F0 mother plants. This resulted in a total of 29 F0 seed families. Seeds were stored under long-term storage conditions, i.e., ultra-dry and at 4°C. Population selection was based on a previous study that elucidated the evolution of cryptic gene pools, the dynamics of the reproductive system, and the diversity of sexual and non-sexual reproductive modes of *Hypericum* in Central Europe, and all populations had been characterized accordingly (Koch *et al*., 2013). On July 17^th^ 2018, these seeds were sown on “Anzucht CLASSIC” peat substrate (https://ökohum.info/aussaat-und-pikiererde/; Ökohum Herbertingen, Germany), and six individuals per F0 seed family were cultivated further in 6 x 6 cm pots and standard “Container CLASSIC” substrate (same provider as peat substrate) mixed with 15% quartz sand. This resulted in a total of 174 F1 individuals. Plants were grown in a climate chamber (16/8 h day/night, 22°C day/18°C night, light conditions 107 µmol/m^2^s^1^) at Heidelberg University. On November 5^th^ 2018, 111 days after sowing, eight leaves per stem from three stems (total 24 leaves) per plant were harvested from the upper part of the plant (starting 4 cm below the tip). In few cases, when only two stems were available or leaves were very small, additional leaves were collected (per stem). Most plants had not yet flowered and no visible infestation by herbivores or infection by pathogens was observed. Leaves were directly shock-frozen in liquid nitrogen, transported to Bielefeld University on dry ice and frozen at -80°C until freeze-drying.

**Table 1.**
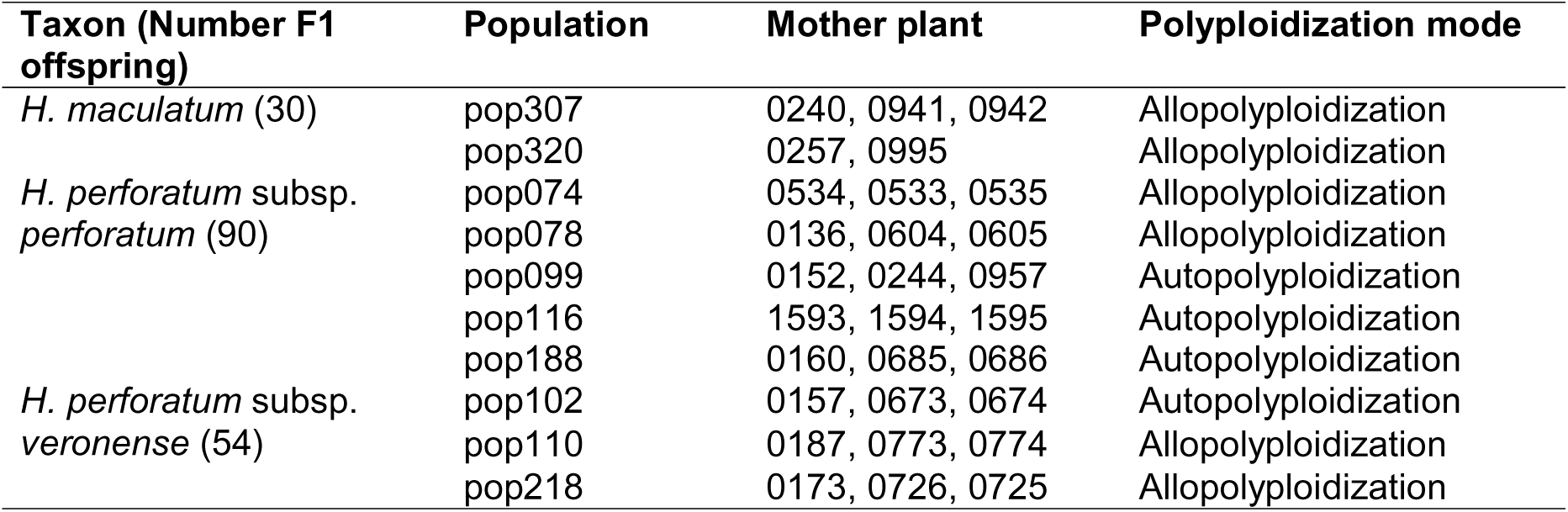
Taxon, population (defined as geographically distinct origin), mother plant, and polyploidization mode of offspring (F1) of each *Hypericum* taxon used for metabolic fingerprinting.

### Genome size analyses and ploidy level assignment

The original individuals from the geographically distinct populations and all F0 mother plants had been studied earlier for chromosome numbers using fresh root tips (Koch *et al*., 2013). In the present study, the respective offsprings (F1 individuals) were studied using genome size estimates for ploidy level assignment relying on the precise F0 karyotypes. Nuclear DNA content was determined using flow cytometry following a simplified protocol (Doležel *et al*., 2007). To release nuclei from the tissue, approximately 50 mm^2^ of fresh and young leaf tissue from each sampled plant was chopped together with approximately 50 mm^2^ of leaf tissue of one internal standard, *Zea mays* cv. CE-777 (2C = 5.43 pg), *Glycine max* cv. Polanka (2C = 2.26 pg) or *Solanum lycopersicum* cv. Stupicke (2C = 1.96 pg), depending on *Hypericum* sample DNA content. The leaf material was homogenized on ice using a sharp razor blade in a Petri dish containing 0.5 mL of ice-cold Nuclei Extraction Buffer from the CyStain^TM^ PI Absolute P kit (ref 05-5022, Sysmec-Partec GmbH, Münster/Görlitz, Germany). The suspension containing the nuclei was filtered through a 30 μm CellTrics® filter (Sysmec-Partec GmbH). Two mL of staining buffer from the CyStain^TM^ PI Absolute P kit, containing propidium iodide and RNase, were added for staining the nuclei and removing the RNA, which could disturb the signal. After 60 min incubation at room temperature in the dark, relative fluorescence intensity of c. 10,000 particles was recorded using a flow cytometer (CyFlow space; Sysmex-Partec GmbH) equipped with a green (532 nm) solid state laser. We applied the following stringent criteria to get precise and stable flow cytometric results: (i) only analyses where the coefficient of variation of the sample peak was below 5% were taken into account, (ii) each sample was measured twice on different days to minimize potential random instrumental drift (Doležel & Bartoš, 2005), and (iii) if the between-day variation exceeded 5%, another measurement was done and the most remote value was discarded. Gating and peak analysis were performed using the Partec FloMax software version 2.4. The 2C peak means of each sample plus the included internal standard were recorded and genome sizes (pg/2C) were calculated from their relative fluorescence intensities.

### Metabolic fingerprinting

Dried leaf material was homogenized and 10 mg were taken for further analysis of (semi-)polar metabolites, following Schweiger et al. (2021), using a subset of *n* = 5 F1 offspring per F0 mother plant plus one offspring for method adjustment. The samples were extracted in 90% ice-cold methanol (*v*:*v*) including hydrocortisone (>98%, Sigma-Aldrich, Steinheim, Germany) as internal standard for 15 min in an ultrasonic bath. They were centrifuged for 10 min at 16,100 g and the supernatants filtered through 0.2 µm filters (Phenomenex, Torrance, CA, USA). The samples and seven blanks were analyzed using an ultra-high performance liquid chromatograph coupled to a quadrupole time-of-flight mass spectrometer (UHPLC-QToF-MS/MS, abbreviated as LC-MS; UHPLC: Dionex UltiMate 3000, Thermo Fisher Scientific, San José, CA, USA; QToF: compact, Bruker Daltonics, Bremen, Germany), equipped with a Kinetex XB-C18 column (150 x 2.1 mm, 1.7 µm, with guard column; Phenomenex) at 45 °C and a flow rate of 0.5 mL /min at a gradient from eluent A, i.e., Millipore-H_2_O with 0.1% formic acid (FA), to eluent B (acetonitrile with 0.1% FA): 2 to 30% B within 20 min, increase to 75 % B within 9 min, followed by column cleaning and equilibration. The QToF was operated in negative electrospray ionization mode and centroid data were taken at a spectra rate of 6 Hz in the *m*/*z* range of 50-1,300. The settings for the MS mode were: end plate offset 500 V, capillary voltage 3,000 V, nebulizer (N_2_) pressure 3 bar, dry gas (N_2_; 275 °C) flow 12 L /min, low mass 90 *m*/*z*, quadrupole ion energy 4 eV, collision energy 7 eV. The AutoMS/MS mode was used for fragmentation (MS/MS), applying isolation widths and collision energies that were increasing along with the precursor *m*/*z*. For recalibration of the *m*/*z* axis, a calibration solution containing sodium formate was introduced into the system prior to each sample. The LC-MS data were processed using the T-ReX 3D algorithm of MetaboScape (v. 2021b, Bruker Daltonics). The processing included *m*/*z* recalibration and picking of metabolic features, each characterized by a specific retention time and *m*/*z* value. Features had to occur in a minimum of 25% of the samples per taxon, with an intensity (peak height) threshold 1,000, and a minimum peak length 20 to be included in the data set. Buckets with metabolic features probably belonging to the same metabolite were created, applying a correlation coefficient threshold of 0.8. Within each bucket, the feature with the highest intensity was used for quantification. Only features in the RT range 1.25-27 min were included, thus removing most primary metabolites co-eluting in the injection peak (<1.2 min). Features were excluded if their intensities were less than 30-fold higher than those in the averaged blanks. The peak heights of the selected features were divided by the peak heights of the hydrocortisone [M+HCOOH-H]^−^ ion (internal standard) and then by the sample dry weights. The feature richness was calculated as the total number of features detected in a sample. Shannon diversity of each sample was calculated from feature richness and evenness based on the relative proportions of the feature intensities. To calculate functional Hill diversity, which considers only features with information on chemical structures, 350 features containing MS/MS data and their corresponding intensities were uploaded to the Global Social Molecular Network of Natural Products (GNPS, https://gnps.ucsd.edu) platform for feature-based molecular networking (FBMN) and calculation of pairwise spectral dissimilarities. Molecular networking was performed with features connected by a similarity score of their MS/MS spectra (parent ion and fragment ion error: 0.02 Da; smallest matching cosine value: 0.7; minimum matched fragment ions: 4). The dissimilarity matrix was extracted from 1-cosine score (parent ion and fragment ion error: 0.02 Da; smallest matching cosine value: 0.01; minimum matched fragment ions: 4). Feature richness, evenness, and dissimilarity matrix were then incorporated into the calculation of functional Hill diversity. The analysis was conducted using the parameter q = 1, meaning that individual features were weighted in the calculation according to their relative proportion (Petrén *et al*., 2023).

### Statistical analyses

All statistical analyses were conducted in R v. 4.5.0 (R Core Team, 2025). Data cleaning, restructuring, and manipulation were performed using *dplyr* (Wickham *et al.,* 2026), *tidyverse* (Wickham *et al.,* 2019), and *janitor* (Firke, 2024).

To visualize variation in metabolic fingerprints, we performed multivariate analyses on the LC-MS dataset. To assess the effects of mother plant and offspring ploidy levels on metabolic fingerprints, we first conducted Partial Least Squares Discriminant Analyses (PLS-DA) for each taxon using *ropls* (Thévenot *et al.,* 2015). For the PLS-DA, data were log_10_-transformed and K-Nearest Neighbor (KNN) imputation of missing values was applied using *VIM* (Kowarik & Templ, 2016), according to Di Guida *et al*. (2016), within each taxon. When predictive models could not be reliably constructed or when predictive accuracy for ploidy level classification was below 0.3, we instead applied non-metric multidimensional scaling (NMDS) using *vegan* (Oksanen *et al.,* 2025). NMDS was based on Bray-Curtis dissimilarities without log_10_-transformation or KNN imputation. Before NMDS analysis, the non-imputed data was subjected to Wisconsin double standardization. Differences in metabolic fingerprints among F0 mother plants, offspring of different ploidy levels, and populations were subsequently tested using permutational multivariate analysis of variance (PERMANOVA) using *vegan* on the corresponding distance matrices.

Shannon diversity, and functional Hill diversity were calculated from the metabolomic feature data using *chemodiv* (Petrén *et al.,* 2023). To evaluate the effects of F0 mother plant and offspring ploidy level on Shannon diversity and functional Hill diversity, we fitted linear mixed effects models (LMMs) using lmer() in *lme4* (Bates *et al.,* 2015) Prior to analysis, Box-Cox transformation using *MASS* (Venables & Ripley, 2002) was applied to response variables that deviated from normality according to Shapiro-Wilk tests. Models were constructed separately for each taxon, with Shannon diversity or functional Hill diversity as the response variables and F0 mother plant and offspring ploidy level as explanatory variables. Owing to sample size limitations, either population or F0 mother plant identity was included as random effect. Models including random effects were compared with fixed-effect-only models using AIC values and likelihood ratio tests, and random effects were retained when they significantly improved model performance. *P-*values of fixed effects were derived using Type II ANOVA in *car* (Fox & Weisberg, 2019). Model diagnostics, including assessments of residual structure and goodness of fit, were performed using *DHARMa* (Hartig, 2024). A similar modelling framework was used to assess the effects of F0 mother plant and offspring ploidy level on feature richness within taxa using generalized linear mixed model fitted with *glmmTMB* (Brooks *et al*., 2017; McGillycuddy *et al.,* 2025) with a negative binomial distribution. Model diagnostics included assessments of overdispersion, zero inflation, and goodness of fit using *DHARMa* (Hartig, 2024). Post hoc pairwise comparisons of estimated marginal means (EMMs) with Tukey adjustment were performed using *emmeans* (Lenth & Piaskowski, 2026) when the overall effect of ploidy level was significant (*p* < 0.05) or marginally significant (0.05 < *p* < 0.10).

Putative molecular formulas, compound names, and chemical classifications, including Natural Product Classification (NPC) pathway, superclass, and class, were annotated using SIRIUS v. 6.1.0 (Dührkop *et al*., 2019, 2021; Ludwig *et al*., 2020) with QToF default settings (10 ppm mass accuracy; de novo plus bottom-up strategy for features below 500 Da). Chemical family was assigned based on superclass only when confidence scores were ≥ 0.8, following recommended criteria (Hoffmann *et al*., 2022, Authier *et al.,* 2026). To visualize variation in major chemical families, heatmaps of total intensities of phenolic acids, flavonoids, terpenoids, and phloroglucinols were generated for each taxon and generation using log_10_-transformed and z-score-standardized feature tables.

Figures were generated primarily using *ggplot2* (Wickham, 2016) and assembled using *patchwork* (Pedersen, 2024). Molecular-network visualizations were produced in Cytoscape v. 3.10.4 (Shannon *et al.,* 2003), and only well-structured clusters with more than five annotated features are presented in the main text. Venn diagrams were generated using *VennDiagram* (Chen & Boutros, 2011). Tables for reporting statistical results were formatted using *flextable* and exported using *officer* (Gohel *et al.,* 2025; Gohel & Skintzos, 2025).

## Results

### Polyploidization is common in all three taxa

In *Hypericum maculatum*, most F0 mother plants were tetraploid, whereas only one plant was hexaploid (Fig. 1A). Among the F1 offspring derived from tetraploid F0 mother plants, most individuals either maintained tetraploidy or increased to pentaploidy, while only a single offspring reached hexaploidy. In contrast, the majority of F1 offspring derived from hexaploid F0 mother plants retained hexaploidy, with only one individual further increasing to heptaploidy. In *H. perforatum* subsp. *perforatum*, the majority of F0 mother plants were tetraploid, whereas only one individual was triploid (Fig. 1B). Offspring derived from the triploid F0 mother plant showed increased ploidy levels, comprising either tetraploid or pentaploid individuals. Among the offspring of tetraploid F0 mother plants, most either remained tetraploid or increased to pentaploidy or hexaploidy. Only two offspring showed reduction from tetraploid to the ancestral diploid state. In *H. perforatum* subsp. *veronense*, F0 mother plants were either diploid or tetraploid (Fig. 1C). F1 offspring derived from diploid F0 plants either retained diploidy or underwent polyploidization, resulting in triploid, tetraploid, or pentaploid offspring. A similar pattern was observed among F1 offspring of tetraploid F0 mother plants, which either maintained tetraploidy or increased to pentaploidy.

**Figure 1.**
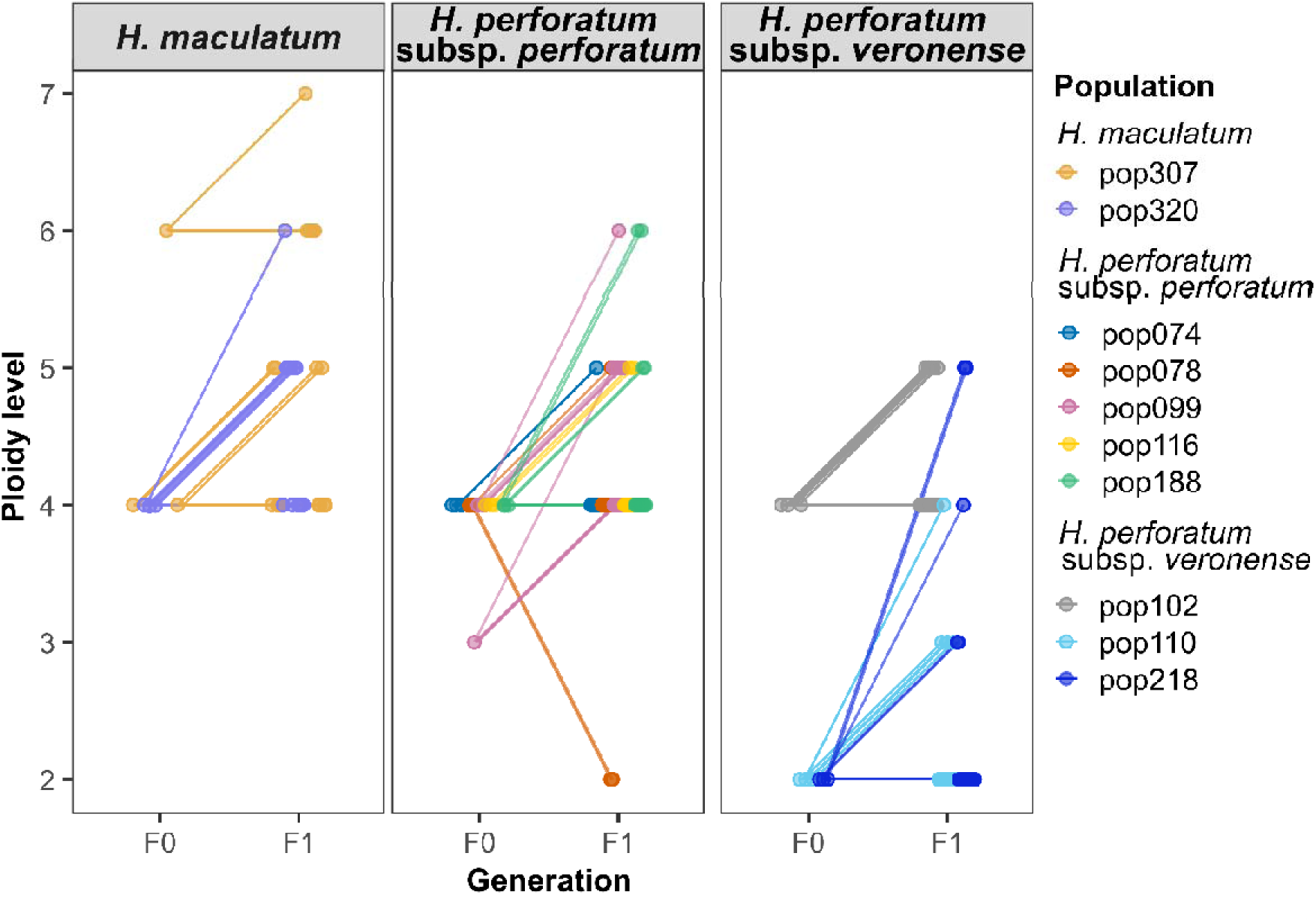
Variation of ploidy levels from F0 to F1 generation across F0 mother plants in *Hypericum* spp. Population codes are provided in Table 1.

### F0 mother plant ploidy levels better discriminated leaf metabolic fingerprints than offspring ploidy levels

Leaf metabolic fingerprints of *H. maculatum* showed a clear group differentiation according to the ploidy level of their F0 mother plants in the score plot of a supervised PLS-DA (Fig. 2A). Metabolic fingerprints differed markedly among F1 offspring derived from tetraploid and hexaploid F0 plants. The PLS-DA model explained 32.5% of the total variance and showed high predictive ability (R^2^X = 0.33, Q^2^ = 0.88), supporting the robustness of F0 mother plant ploidy level as a classifier of leaf metabolic fingerprints. In contrast, PLS-DA model construction failed based on F1 ploidy level (data not shown). Therefore, the effect of F1 ploidy level was further evaluated using unsupervised NMDS ordination and PERMANOVA. PERMANOVA results indicated that both F1 ploidy level and population significantly affected the metabolic fingerprints (*df* = 3, *F* = 1.59, *p* = 0.007 and *df* = 3, *F* = 4.73, *p* < 0.001, respectively). Consistent with these results, NMDS ordination revealed a stronger clustering by population than by F1 ploidy level, with only limited separation among ploidy levels (Fig. 2B).

**Figure 2.**
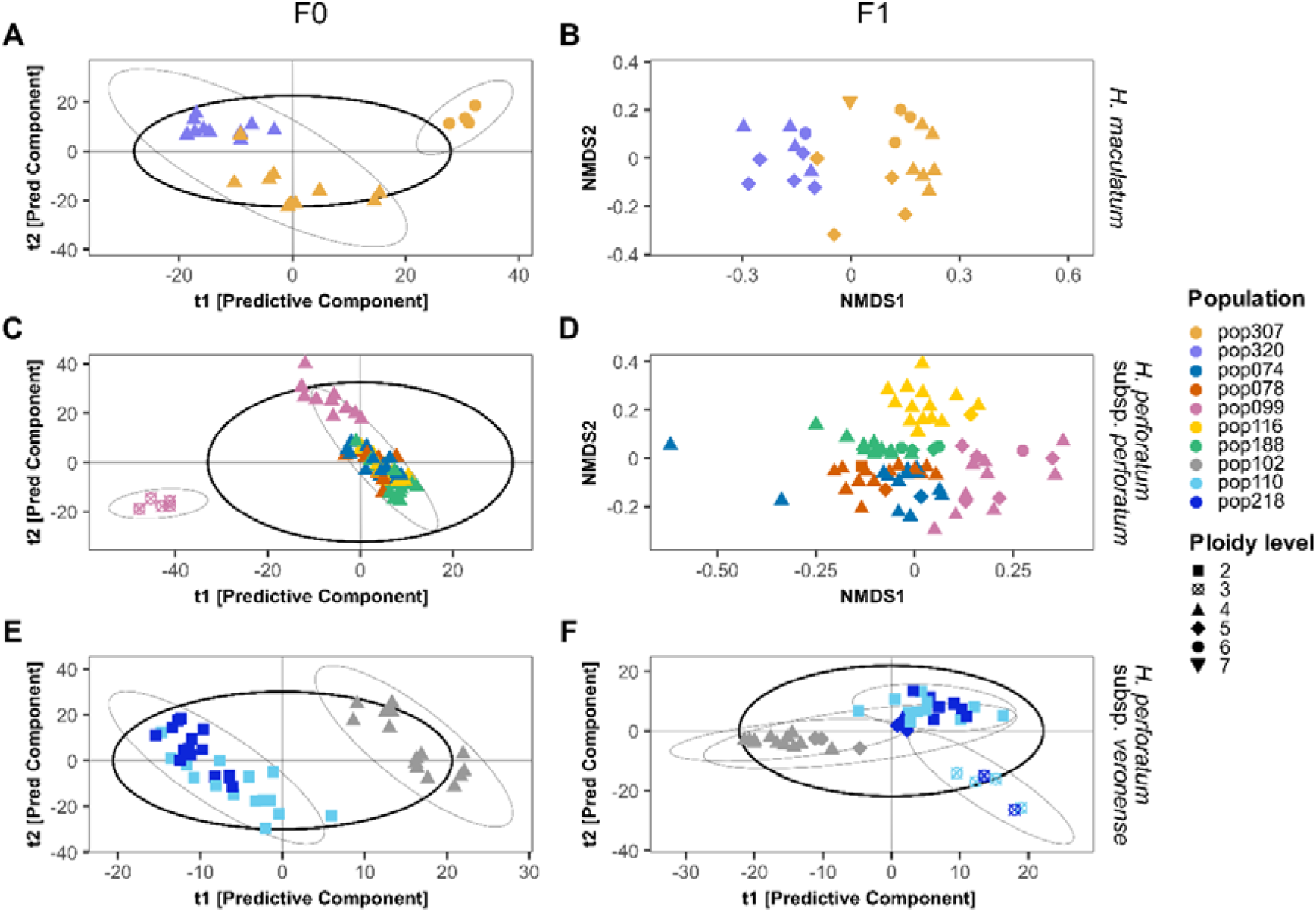
Partial least squares discriminant analyses (PLS-DA; A, C, E, F) or non-metric multidimensional scaling (NMDS; B, D) of leaf metabolic fingerprints of offspring individuals (F1) of *Hypericum* spp. grown under common climate chamber conditions. *Hypericum maculatum* (A, B; based on 2,292 metabolic features), *Hypericum perforatum* subsp. *perforatum* (C, D; based on 2,336 features), and *Hypericum perforatum* subsp. *veronense* (E, F; based on 2,288 features). F0 indicates ploidy level of F0 mother plant, and F1 indicates ploidy level of F1 offspring. Symbols represent ploidy level, while colors indicate population identity. Thick black and thin grey ellipses represent overall and ploidy level-specific 95% confidence intervals, respectively.

A similar pattern was observed in *H. perforatum* subsp. *perforatum*. F0 mother plant ploidy level explained variation in the metabolic fingerprints of F1 offspring (Fig. 2C), with distinct separation between F1 offspring derived from triploid and tetraploid F0 plants (PLS-DA: R^2^X = 0.21, Q^2^ = 0.94). By contrast, F1 ploidy level did not support a reliable supervised classification, as PLS-DA model construction failed (data not shown). We therefore assessed the impact of F1 ploidy level using PERMANOVA. F1 ploidy had no significant effect on metabolic fingerprints (PERMANOVA: *df* = 3, *F* = 1.18, *p* = 0.13), whereas population had a significant effect (*df* = 3, *F* = 5.67, *p* < 0.001). Consistently, NMDS further indicated that metabolic fingerprints were rather structured by population than by F1 ploidy level (Fig. 2D).

In *H. perforatum* subsp. *veronense*, both F0 mother plant ploidy level and F1 ploidy level influenced the metabolic fingerprints of F1 offspring (Fig. 2E, F). However, the effect of the F0 mother plant ploidy level was more pronounced than that of the F1 ploidy level (PLS-DA; F0 mother plant ploidy level: R^2^X = 0.26, Q^2^ = 0.92; F1 ploidy level: R^2^X = 0.11, Q^2^ = 0.28). F1 offspring derived from diploid and tetraploid F0 plants formed distinct groups irrespective of population origin (Fig. 2E). In contrast, separation among F1 ploidy levels was weaker, with pentaploid individuals showing substantial overlap with both diploid and tetraploid F1 plants (Fig. 2F).

### Ploidy increase is partly associated with a gain of chemical features

In total, 2,485 metabolic features were detected in the leaves of all three *Hypericum* taxa. In *H. maculatum,* 2,292 features were detected (Fig. 3A), including a large core set (1,926 features) shared among all F1 ploidy levels. Individuals of each ploidy level also contained unique features, with pentaploid plants harboring the highest number of unique features. In *H. perforatum* subsp. *perforatum*, 2,336 features were detected (Fig. 3B), of which 1,973 features were shared among individuals of all ploidy levels. Diploid plants were the ones without unique metabolic features, whereas individuals of all the other polyploidy levels possessed distinct subsets of unique features. Tetraploid plants showed the most unique features, but feature number did not consistently increase with ploidy level. Similarly, in *H. perforatum* subsp. *veronense*, 2,288 features were detected, of which 2,027 were shared across individuals of all ploidy levels (Fig. 3C). Compared to individuals of all other ploidy levels, triploid plants exhibited the highest feature number with 82 additional features. However, feature number did not consistently increase with ploidy level.

**Figure 3.**
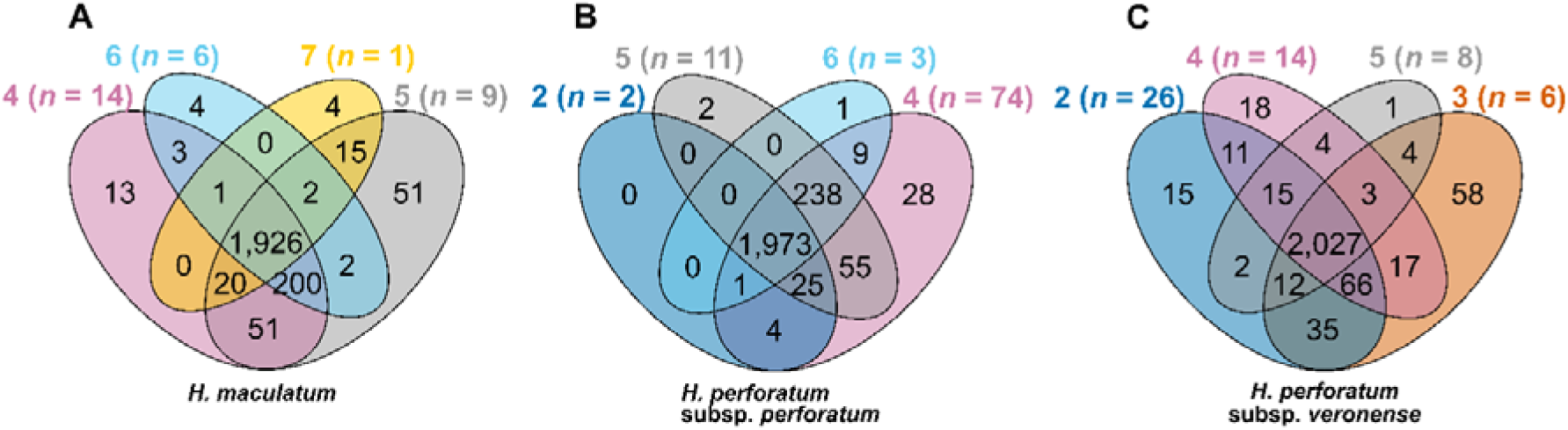
Venn diagrams showing the overlap of metabolic features in leaves across offspring individuals (F1) of different ploidy levels in *Hypericum* ssp. (A) *Hypericum maculatum*; (B) *Hypericum perforatum* subsp. *perforatum*; (C*) Hypericum perforatum* subsp. *veronense*. The bold numbers on the top and the different colors indicate distinct ploidy levels, while numbers in parentheses represent the sample size for each ploidy level.

### Ploidy levels affect leaf chemodiversity

In *H. maculatum*, the Shannon diversity of the metabolic features in leaves of F1 plants tended to decline with increasing F0 mother plant ploidy level, with lower diversity in offspring from hexaploid than in offspring from tetraploid F0 mother plants (Table 2; Fig. 4A). In *H. perforatum* subsp. *perforatum*, Shannon diversity, functional Hill diversity, and feature richness were all influenced by the ploidy level of either the F0 mother plants and/ or the F1 offspring, although the strength of these effects varied (Table 2; Fig. 4B-E). Both Shannon diversity and functional Hill diversity increased with F0 mother plant ploidy level, with higher diversity in offspring from tetraploid than in offspring from triploid F0 mother plants (Fig. 4B, C). In contrast, functional Hill diversity and feature richness were marginally associated with F1 ploidy level; however, effects were not reflected in pairwise comparisons among ploidy levels (Table 2; Fig. 4D, E). In *H. perforatum* subsp. *veronense*, only the feature richness was significantly affected by ploidy level (Table 2), with lower feature richness in offspring from diploid than in offspring from tetraploid F0 mother plants. Among F1 ploidy levels, triploid offspring showed higher feature richness than both diploid and tetraploid offspring (Fig. 4F, G). Neither Shannon diversity nor functional Hill diversity were significantly associated with ploidy level in this taxon (Table 2).

**Figure 4.**
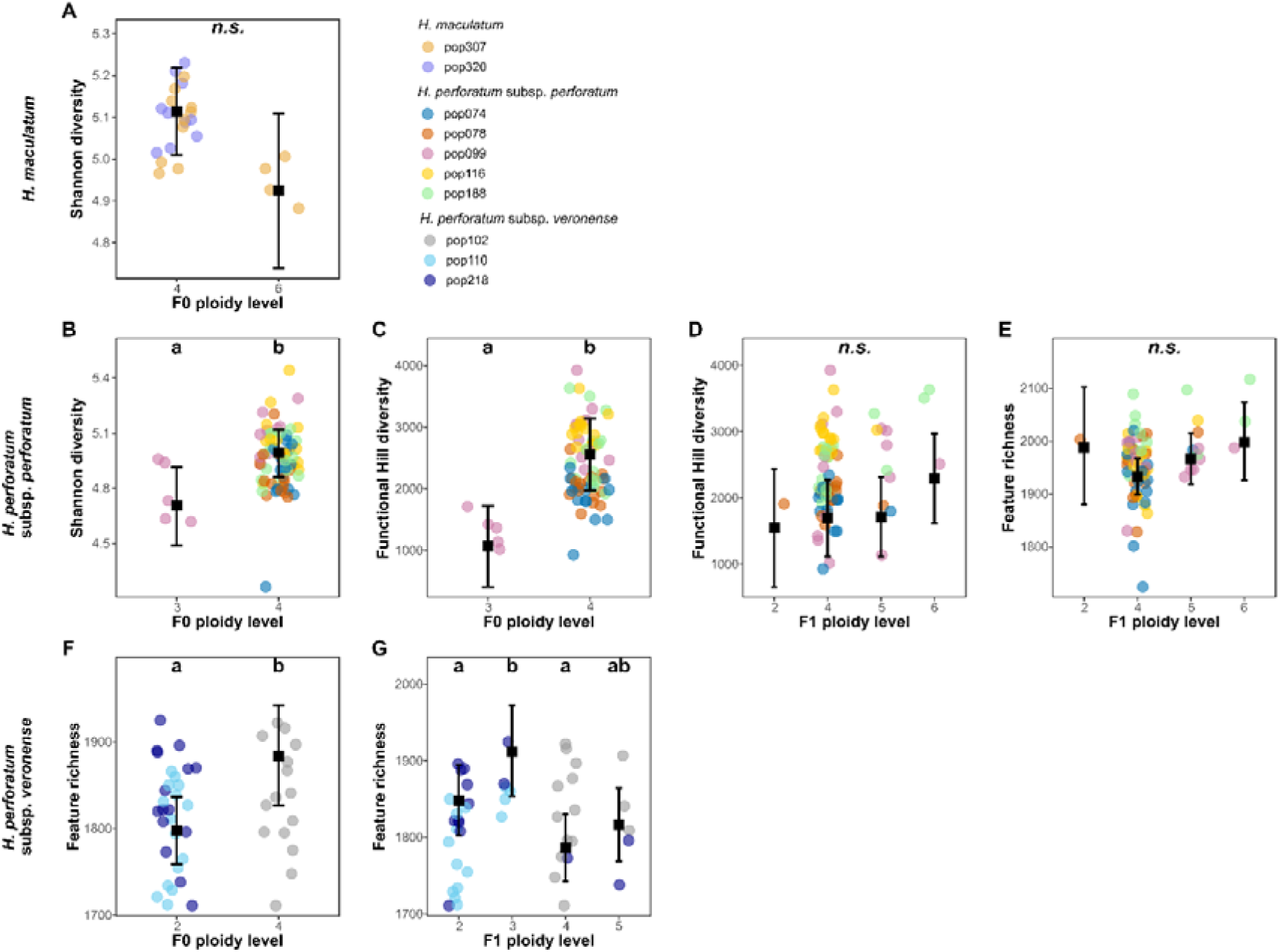
Shannon diversity, functional Hill diversity, and richness of leaf metabolic features of F1 offspring of *Hypericum*: *Hypericum maculatum* (A), *H. perforatum* subsp. *perforatum* (B-E), and *H. perforatum* subsp. *veronense* (F, G). F0 indicates F0 mother plant, and F1 indicates F1 offspring. Black error bars indicate estimated marginal means (EMMs) with 95% confidence intervals, and the black squares indicate the corresponding EMM estimates. Individual observations are shown as circles. Only response variables that were (marginally) significantly affected by ploidy levels (see Table 1) are depicted. Different letters indicate significant pairwise differences among ploidy levels based on EMM post hoc comparisons with Tukey adjustment (*p* < 0.05); *n.s.*, not significant.

**Table 2.**
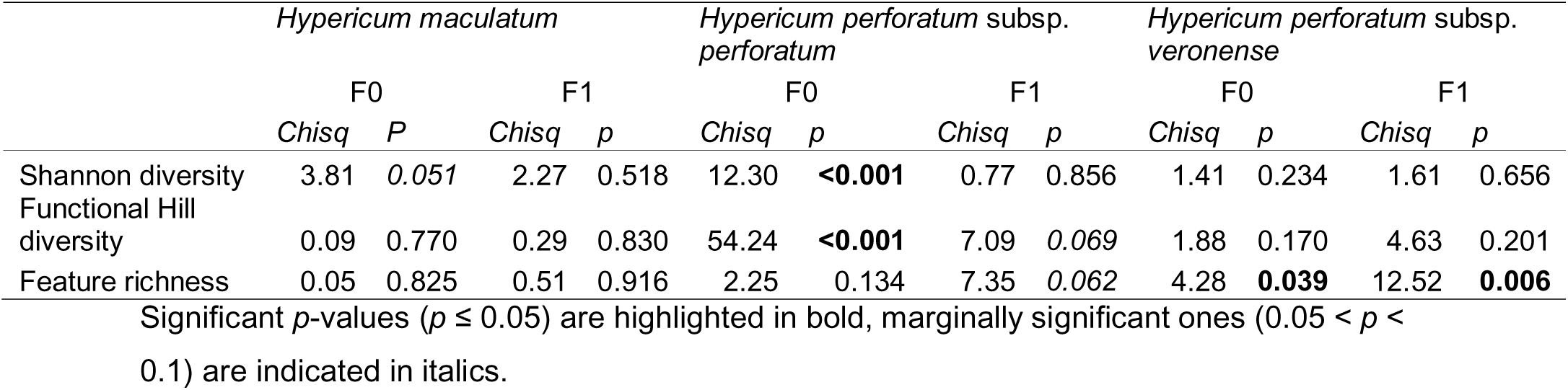
Summary of *Chisq* and *p*-values from ANOVAs based on (generalized) linear (mixed effects) models, testing the effects of F0 ploidy level and F1 ploidy level on Shannon diversity, functional Hill diversity, and feature richness of leaf metabolic fingerprints of offspring individuals (F1) of *Hypericum* spp.

|  | *Hypericum maculatum* |  |  |  | *Hypericum perforatum* subsp. *perforatum* |  |  |  | *Hypericum perforatum* subsp. *veronense* |  |  |  |
| --- | --- | --- | --- | --- | --- | --- | --- | --- | --- | --- | --- | --- |
|  | F0 |  | F1 |  | F0 |  | F1 |  | F0 |  | F1 |  |
|  | *Chisq* | *P* | *Chisq* | *p* | *Chisq* | *p* | *Chisq* | *p* | *Chisq* | *p* | *Chisq* | *p* |
| Shannon diversity | 3.81 | 0.051 | 2.27 | 0.518 | 12.30 | **<0.001** | 0.77 | 0.856 | 1.41 | 0.234 | 1.61 | 0.656 |
| Functional Hill diversity | 0.09 | 0.770 | 0.29 | 0.830 | 54.24 | **<0.001** | 7.09 | 0.069 | 1.88 | 0.170 | 4.63 | 0.201 |
| Feature richness | 0.05 | 0.825 | 0.51 | 0.916 | 2.25 | 0.134 | 7.35 | 0.062 | 4.28 | **0.039** | 12.52 | **0.006** |

Significant *p*-values ( $p \leq 0.05$ ) are highlighted in bold, marginally significant ones ( $0.05 < p < 0.1$ ) are indicated in italics.

### Intensities of specific chemical families vary across ploidy levels

Feature-based molecular networking yielded 16 distinct molecular clusters (Fig. S2). Of these, three clusters were dominated by phenolic acids, flavonoids, terpenoids, and phloroglucinols, while a substantial proportion of nodes remained unannotated (Fig. 5). In *H. maculatum*, total intensities (sum of intensities of features within a chemical family) within these annotated chemical families were generally lower in F1 offspring derived from hexaploid F0 mother plants than in F1 offspring derived from tetraploid F0 mother plants.

**Figure 5.**
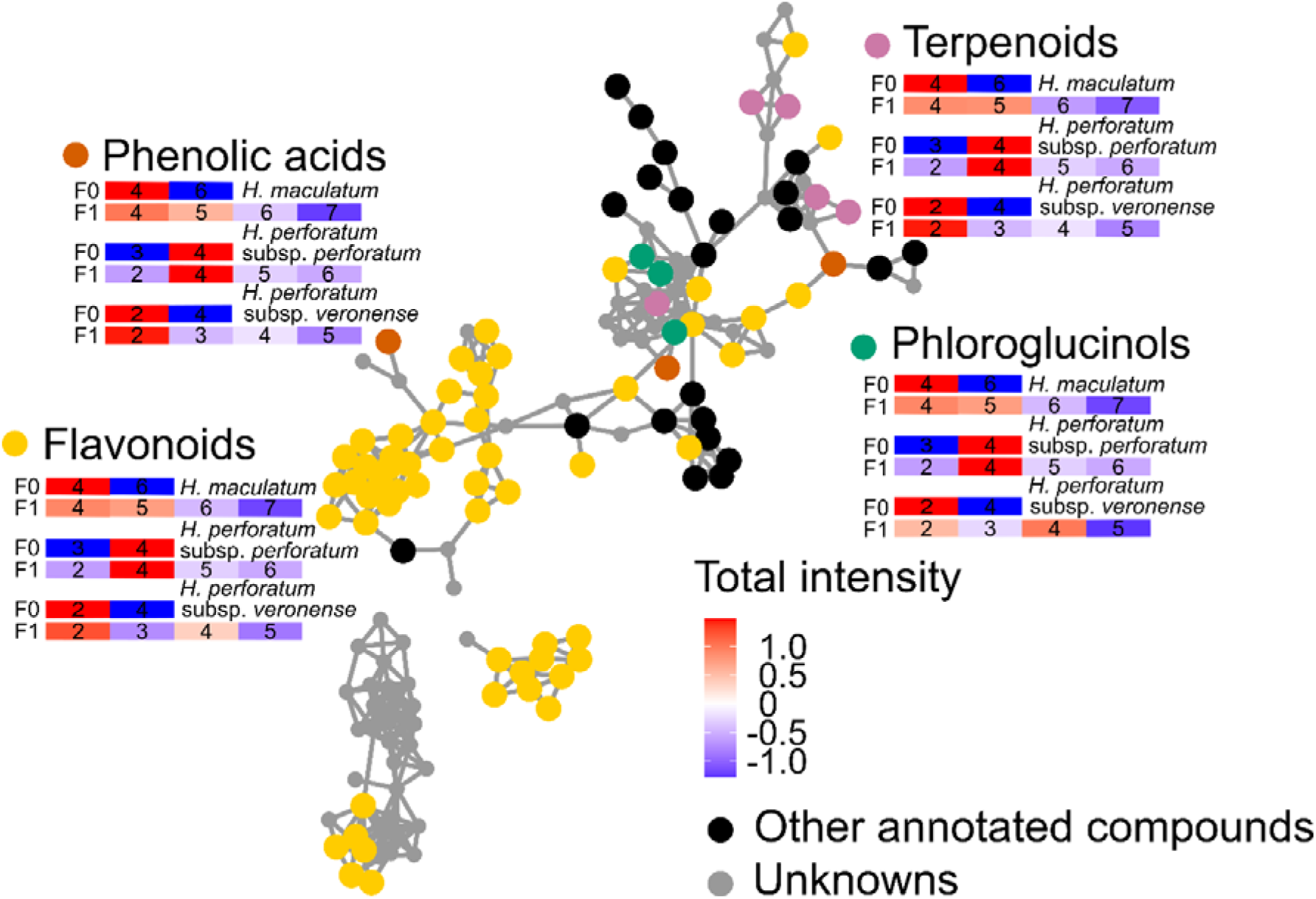
Scaled variation in chemical families of leaf metabolic features in offspring individuals (F1) of three *Hypericum* taxa with ploidy levels of F0 mother plants and of F1 offspring, based on a feature-based molecular network constructed from 350 features measured via LC-MS. F0 indicates the ploidy level of the F0 mother plant, while F1 indicates the ploidy level of F1 offspring. Features (nodes), identified using SIRIUS (with confidence scores ≥ 0.8), were colored according to chemical families. Edges represent similarities in MS/MS fragmentation patterns, providing a visual measure of structural relatedness among features. Only well-structured clusters with more than five annotated features are presented. Ploidy levels are indicated within each rectangle. The total intensity of each chemical family was calculated based on all annotated features within this family.

Across F1 offspring, total intensities of all above mentioned chemical families declined in individuals from tetraploid to heptaploid levels. In contrast, in *H. perforatum* subsp. *perforatum*, feature intensities increased with F0 ploidy level. F1 offspring derived from triploid F0 mother plants generally showed lower total intensities than offspring derived from tetraploid F0 mother plants. Among F1 individuals, feature intensities increased from diploid to tetraploid individuals but did not continue to rise in pentaploid and hexaploid plants.

However, similar to *H. maculatum*, in *H. perforatum* subsp. *veronense* total feature intensities were lower in F1 offspring derived from F0 tetraploid plants than F1 offspring derived from diploid F0 mother plants. Among F1 offspring individuals, diploids exhibited the highest feature intensities, followed by tetraploids, whereas triploids and pentaploids consistently showed the lowest intensities in phenolic acids, flavonoids, and terpenoids.

## Discussion

Polyploidization is widely recognized as an important evolutionary force in plant diversification and speciation and may also have substantial effects on chemodiversity (Wood *et al*., 2009). Although several hypotheses have been proposed to explain the drivers and evolution of chemodiversity (Wetzel & Whitehead, 2020; Thon *et al*., 2024), the role of polyploidization in shaping metabolite diversity has remained comparatively underexplored. To address this gap, we assessed ploidy levels in plants of *Hypericum maculatum* and *H. perforatum* ssp. sampled from ten populations distributed across Europe and grown under common climate chamber conditions. We then compared leaf metabolic fingerprints, Shannon diversity, functional Hill diversity, feature richness as well as intensities of key chemical families identified during our analysis in relation to mother plant and offspring ploidy levels. We found that ploidy level had a pronounced effect on the metabolic fingerprints, Shannon diversity, functional Hill diversity, and feature richness. However, the strength and direction of these effects varied both between and within taxa.

Polyploidization is a recurrent and fundamental evolutionary process in *Hypericum* and its high occurrence is likely related to a mixed reproductive system combining sexual reproduction with varying degrees of facultative pseudogamous apomixis (Koch *et al*., 2013). Apomixis can stabilize novel genomic configurations by bypassing meiotic constraints, thereby promoting the establishment of polyploid lineages that might otherwise suffer reduced fertility or genomic instability following whole-genome duplication (Hojsgaard, 2018). High rates of polyploidy formation have also been reported in other facultatively apomictic species, such as *Ranunculus carpaticola* (Ranunculaceae) and *Paspalum notatum* (Poaceae) (Paun *et al*., 2006; Escobar *et al*., 2025). Notably, in addition to polyploid events, haploid parthenogenesis has been observed in *H. perforatum* in this study, indicating that a facultative apomictic reproductive system may facilitate the persistence of rare ploidy events.

Contrary to our hypothesis, the ploidy level of F0 mother plants explained more variation in leaf metabolic fingerprints of F1 offspring than did the F1 offspring ploidy level. During sexual reproduction, chromosome segregation and recombination generate novel allelic combinations that can disrupt dosage-sensitive regulatory networks, potentially leading to widespread changes in gene expression and metabolite accumulation (Mason & Wendel, 2020). In contrast, apomictic reproduction may stabilize such altered metabolic states by maintaining clonal inheritance of an initial genomic configuration. However, this stabilizing effect may be counteracted by polyploidization via subsequent regulatory adjustments and polyploidy-associated epigenetic silencing, which may lead to metabolic divergence among offspring lineages of different ploidy levels (Adams & Wendel, 2005; Xu *et al*., 2022).

Offspring of different ploidy levels in *Hypericum* spp. is a result of both sexual and apomictic reproduction, which may act together with epigenetic modifications linked to polyploidization. Together, these interacting mechanisms may explain the reduced discriminatory power of F1 offspring ploidy level for leaf metabolic fingerprints relative to that of F0 mother plants across all three *Hypericum* taxa.

Partially aligning with our hypothesis, the consequences of polyploidization on metabolic fingerprints, on the presence of metabolic features, and on feature intensities were non-linear and context-dependent across the three *Hypericum* taxa studied herein. Polyploidization resulted in both unique features and shifts in the intensity of major chemical families, but responses were inconsistent and not predictable from ploidy level alone. Notably, even low-ploidy F1 offspring in *H. maculatum* and *H. perforatum* subsp. *veronense* produced unique features absent from individuals with higher polyploidy levels. This finding suggests that substantial metabolic modification can arise through the segregation and recombination of polyploid F0 mother plants, rather than through changes in ploidy levels alone. This is consistent with findings in *Arabidopsis thaliana* (Brassicaceae), where unique features were detected in both parents and offspring through inbreeding (Keurentjes *et al*., 2006). At the same time, the present study revealed lower intensities of major chemical families in lineages of higher F0 ploidy levels in *H*. *maculatum* and *H. perforatum* subsp. *veronense* compared to lineages with the lowest ploidy levels, represented by tetraploid or diploid plants, respectively. This finding aligns with previous reports of lower hypericin contents (a phloroglucinol derivative) in tetraploid *H. perforatum* compared to diploids (Čellárová *et al*., 1997). Such reduction may reflect a dilution effect following genome duplication, as demonstrated for metabolite abundance per cell in *Spirodela polyrhiza* (Araceae) (Wu *et al*., 2024). In contrast, tetraploid F1 individuals of *H. perforatum* subsp. *perforatum* showed the highest feature intensities across all tested chemical families compared with individuals of lower or higher ploidy in the present study, highlighting the absence of a universal direction of response. Overall, these findings demonstrate that the effects of polyploidization on plant metabolic fingerprints and intensities of chemical families are shaped by interactions among ploidy level, genetic background, and regulatory processes.

Beyond effects on the overall leaf metabolic fingerprints and intensities of different chemical families, polyploidization also influenced chemodiversity in all three taxa. However, only partly supporting our hypothesis, the responses varied among diversity metrics and were taxon-specific. In *H. maculatum*, Shannon diversity decreased with increasing F0 mother plant ploidy level, potentially reflecting epigenetic silencing-associated reconfiguration of metabolic pathways. Such polyploidization-induced shifts in primary and specialized metabolite profiles have been reported in *Hylocereus* spp. (Cactaceae), although the direction and magnitude of these responses were highly taxon- and metabolite-specific (Cohen *et al*., 2013). In contrast, functional Hill diversity increased with F0 mother plant ploidy level in *H. perforatum* subsp. *perforatum* in the present study, suggesting that polyploidization may enhance the expression of otherwise weakly active biosynthetic pathways and promote structural diversification of metabolites. This finding is in line with qualitative and quantitate changes in specialized metabolite profiles reported across other medicinal plants (Madani *et al*., 2021).

Although both *H. perforatum* subsp. *perforatum* and *H. perforatum* subsp. *veronense* include allopolyploidization and autopolyploidization populations, the mode of polyploidization did not show obvious shifts in chemodiversity across individuals differing in F0 or F1 ploidy levels. This contrasts with the expectation that allopolyploidization should generate stronger metabolic novelty than autopolyploidization (Soltis *et al.,* 2014). Instead, our results suggest that polyploidization per se, rather than the specific mode of polyploidization, may be more relevant for shaping chemodiversity in *Hypericum*. This interpretation is consistent with evidence that autopolyploidization alone can remodel plant metabolomes, although the direction and magnitude of these effects are often lineage- and compound-specific (Fasano *et al*., 2016).

Notably, F0 mother plant ploidy level explained chemodiversity patterns better than offspring ploidy level in our *Hypericum* data, suggesting that evolutionary history can outweigh the immediate consequences of genome duplication. Despite extensive work on the influence of polyploidization on feature intensities and metabolic fingerprints (Gaynor *et al*., 2020), impacts on chemodiversity have received little attention, although chemodiversity is increasingly recognized as an important component of plant ecological interactions and evolution (Wood *et al*., 2009; Wetzel & Whitehead, 2020). Our results suggest that polyploidization may represent one rapid mechanism contributing to the evolution of plant chemodiversity. This may be particularly relevant in *H. perforatum*, which has successfully invaded temperate regions worldwide and is used as a medical plant. The high frequency of polyploidization and the associated swift divergence in chemodiversity in *H. perforatum* may contribute to its ecological success and adaptive potential, while also providing opportunities for the discovery of novel bioactive compounds. However, ploidy levels were unevenly represented in our dataset, and not all polyploid lineages were sampled across the studied taxa, limiting our ability to draw general conclusions regarding the relationship between ploidy and chemodiversity. Future studies should therefore include a broader range of taxa and ploidy levels to fully understand the role of polyploidization in shaping plant chemodiversity.

## Acknowledgement

We thank Susanne Hoibian for assistance in LC-MS analysis and Peter Sack for genome size estimates. We are also grateful to the gardeners at Heidelberg Botanical Garden for plant care and cultivation. This study was inspired by the research unit FOR 3000 (project number 415496540), funded by the Deutsche Forschungsgemeinschaft (DFG) (MU1829/28-2).

## Author contributions

MAK and CM planned and designed the research. MAK sampled and grew the plants. CM harvested the experimental plants, RS and CM performed the metabolic fingerprinting analyses, and ERS and MAK performed the ploidy level analyses. XX analyzed the data and wrote the initial draft of the manuscript with input from RS, ERS, TD, MAK, and CM. All authors interpreted the data and contributed to the final version of this paper.

## Data availability

The datasets used and/or analyzed during the current study are available from the corresponding author and will be made available via MetaboLights (Yurekten *et al*., 2024) and DataPlant upon publication of this manuscript.

## References

Adams KL, Wendel JF. 2005. Polyploidy and genome evolution in plants. Current Opinion in Plant Biology 8: 135–141.

Authier E, Frachon L, Friedrichs J, Brokate L, Junker RR, Müller C, Dussarrat T. 2026. Metabolism and chemical diversity evolve in response to pollinator availability. bioRxiv DOI: 10.64898/2026.02.13.702789

Barnes J, Anderson LA, Philipson JD. 2001. St John’s wort (*Hypericum perforatum* L.): a review of its chemistry, pharmacology and clinical properties. Journal of Pharmacy and Pharmacology 53: 583–600.

Bates D, Mächler M, Bolker BM, Walker SC. 2015. Fitting linear mixed-effects models using lme4. Journal of Statistical Software 67: 1–48.

Brooks ME, Kristensen K, van Benthem KJ, Magnusson A, Berg CW, Nielsen A, Skaug HJ, Maechler M, Bolker BM. 2017. glmmTMB Balances Speed and Flexibility Among Packages for Zero-inflated Generalized Linear Mixed Modeling. The R Journal 9: 378–400.

Čellárová E, Brutovská R, Bruňáková K, Daxnerová Z, Weigel RC. 1997. Correlation between hypericin content and the ploidy of somaclones of *Hypericum perforatum* L. Acta Biotechnologica 17: 8390.

Chen H, Boutros PC. 2011. VennDiagram: a package for the generation of highly-customizable Venn and Euler diagrams in R. BMC Bioinformatics 12: 35.

Cohen H, Fait A, Tel-Zur N. 2013. Morphological, cytological and metabolic consequences of autopolyploidization in *Hylocereus* (Cactaceae) species. BMC Plant Biology 13: 173.

Di Guida R, Engel J, Allwood JW, Weber RJM, Jones MR, Sommer U, Viant MR, Dunn WB. 2016. Non-targeted UHPLC-MS metabolomic data processing methods: a comparative investigation of normalisation, missing value imputation, transformation and scaling. Metabolomics 12: 93.

Doležel J, Bartoš J. 2005. Plant DNA flow cytometry and estimation of nuclear genome size. Annals of Botany 95: 99–110.

Doležel J, Greilhuber J, Suda J. 2007. Estimation of nuclear DNA content in plants using flow cytometry. Nature Protocols 2: 2233–2244.

Dührkop K, Fleischauer M, Ludwig M, Aksenov AA, Melnik AV, Meusel M, Dorrestein PC, Rousu J, Böcker S. 2019. SIRIUS 4: a rapid tool for turning tandem mass spectra into metabolite structure information. Nature Methods 16: 299–302.

Dührkop K, Nothias L-F, Fleischauer M, Reher R, Ludwig M, Hoffmann MA, Petras D, Gerwick WH, Rousu J, Dorrestein PC et al. 2021. Systematic classification of unknown metabolites using high-resolution fragmentation mass spectra. Nature Biotechnology 39: 462–471.

Escobar LM, Reutemann AV, Perichon MC, Schneider JS, Sartor CA, Chaparro C, Daviña JR, Valls JFM, Martínez EJ, Honfi AI. 2025. Neonative Diploid-Polyploid Hotspots of *Paspalum notatum*: Identifying Novel Genetic Diversity for Conservation in South America. Genes 16: 1098.

Fasano C, Diretto G, Aversano R, D’Agostino N, Di Matteo A, Frusciante L, Giuliano G, Carputo D. 2016. Transcriptome and metabolome of synthetic *Solanum* autotetraploids reveal key genomic stress events following polyploidization. New Phytologist 210: 1382–1394.

Firke S. 2024. janitor: Simple Tools for Examining and Cleaning Dirty Data. [WWW document] URL https://CRAN.R-project.org/package=janitor [accessed 23 May 2026].

Fox J, Weisberg S. 2019. An R Companion to Applied Regression. Thousand Oaks CA: Sage.

Gaynor ML, Lim-Hing S, Mason CM. 2020. Impact of genome duplication on secondary metabolite composition in non-cultivated species: a systematic meta-analysis. Annals of Botany 126: 363–376.

Ghasemi Pirbalouti A, Fatahi-Vanani M, Craker L, Shirmardi H. 2014. Chemical composition and bioactivity of essential oils of *Hypericum helianthemoides*, *Hypericum perforatum* and *Hypericum scabrum*. Pharmaceutical Biology 52: 175–181.

Gohel D, Moog S, Heckmann M. 2025. officer: Manipulation of Microsoft Word and PowerPoint Documents. [WWW document] URL https://CRAN.R-project.org/package=officer [accessed 23 May 2026].

Gohel D, Skintzos P. 2025. flextable: Functions for Tabular Reporting. [WWW document] URL https://CRAN.R-project.org/package=flextable [accessed 23 May 2026].

Hanusch M, Dötterl S, Larue-Kontić A-AC, Keller A, Junker RR. 2025. Floral scent chemodiversity is associated with high floral visitor but low bacterial richness on flowers. New Phytologist 248: 3270–3279.

Hartig F. 2024. DHARMa: Residual Diagnostics for Hierarchical (Multi-Level / Mixed) Regression Models. [WWW document] URL https://CRAN.R-project.org/package=DHARMa [accessed 23 May 2026].

Hoffmann MA, Nothias L-F, Ludwig M, Fleischauer M, Gentry EC, Witting M, Dorrestein PC, Dührkop K, Böcker S. 2022. High-confidence structural annotation of metabolites absent from spectral libraries. Nature Biotechnology 40: 411–421.

Hojsgaard D. 2018. Transient Activation of Apomixis in Sexual Neotriploids May Retain Genomically Altered States and Enhance Polyploid Establishment. Frontiers in Plant Science 9: 230.

Jesus-Gonzalez L de, Weathers PJ. 2003. Tetraploid *Artemisia annua* hairy roots produce more artemisinin than diploids. Plant Cell Reports 21: 809–813.

Keurentjes JJB, Fu J, Vos CHR de, Lommen A, Hall RD, Bino RJ, van der Plas LHW, Jansen RC, Vreugdenhil D, Koornneef M. 2006. The genetics of plant metabolism. Nature Genetics 38: 842–849.

Koch MA, Scheriau C, Betzin A, Hohmann N, Sharbel TF. 2013. Evolution of cryptic gene pools in *Hypericum perforatum*: the influence of reproductive system and gene flow. Annals of Botany 111: 1083–1094.

Koperdáková J, Kosuth J, Cellárová E. 2007. Variation in the content of hypericins in four generations of seed progeny of *Hypericum perforatum* somaclones. Journal of Plant Research 120: 123–128.

Kowarik A, Templ M. 2016. Imputation with the R Package VIM. Journal of Statistical Software 74: 1–16.

Lenth R, Piaskowski J. 2026. emmeans: Estimated Marginal Means, aka Least-Squares Means. [WWW document] URL https://rvlenth.github.io/emmeans/. [accessed 10 August 2026]

Lichman BR, Godden GT, Buell CR. 2020. Gene and genome duplications in the evolution of chemodiversity: perspectives from studies of Lamiaceae. Current Opinion in Plant Biology 55: 74–83.

Ludwig M, Nothias L-F, Dührkop K, Koester I, Fleischauer M, Hoffmann MA, Petras D, Vargas F, Morsy M, Aluwihare L et al. 2020. Database-independent molecular formula annotation using Gibbs sampling through ZODIAC. Nature Machine Intelligence 2: 629–641.

Madani H, Escrich A, Hosseini B, Sanchez-Muñoz R, Khojasteh A, Palazon J. 2021. Effect of Polyploidy Induction on Natural Metabolite Production in Medicinal Plants. Biomolecules 11: 899.

Malacrinò A, Jakobs R, Xu S, Müller C. 2025. Influences of plant maternal effects, chemotype, and environment on the leaf bacterial community. Plant Biology 27: 903–912.

Manteghi Tafreshi A, Mohammadhassan R. 2025. Sodium chloride, colchicine, and 6-benzylaminopurine can change antioxidant property and phenols content in *Hypericum perforatum* L.: An in vitro study. Acta Agriculturae Slovenica 121/3: 1–12.

Mason AS, Wendel JF. 2020. Homoeologous Exchanges, Segmental Allopolyploidy, and Polyploid Genome Evolution. Frontiers in Genetics 11: 1014.

McGillycuddy M, Warton DI, Popovic G, Bolker BM. 2025. Parsimoniously Fitting Large Multivariate Random Effects in glmmTMB. Journal of Statistical Software 112: 1–19.

Molins MP, Corral JM, Aliyu OM, Koch MA, Betzin A, Maron JL, Sharbel TF. 2014. Biogeographic variation in genetic variability, apomixis expression and ploidy of St. John’s wort (*Hypericum perforatum*) across its native and introduced range. Annals of Botany 113: 417–427.

Müller C, Junker RR. 2022. Chemical phenotype as important and dynamic niche dimension of plants. New Phytologist 234: 1168–1174.

Ojeda-Prieto L, Moreno EL, Heinen R, Weisser WW. 2025. Intraspecific plant chemodiversity at plot level has contrasting effects on arthropod functional groups. Functional Ecology 39: 3732–3750.

Oksanen J, Simpson GL, Blanchet FG, Kindt R, Legendre P, Minchin PR, O’Hara RB, Solymos P. 2025. vegan: Community Ecology Package. [WWW document] URL https://CRAN.R-project.org/package=vegan [accessed 23 May 2026].

Paun O, Greilhuber J, Temsch EM, Hörandl E. 2006. Patterns, sources and ecological implications of clonal diversity in apomictic *Ranunculus carpaticola* (*Ranunculus auricomus* complex, Ranunculaceae). Molecular Ecology 15: 897–910.

Pedersen TL. 2024. patchwork: The Composer of Plots. [WWW document] URL https://CRAN.R-project.org/package=patchwork.

Petrén H, Anaia RA, Aragam KS, Bräutigam A, Eckert S, Heinen R, Jakobs R, Ojeda-Prieto L, Popp M, Sasidharan R et al. 2024. Understanding the chemodiversity of plants: Quantification, variation and ecological function. Ecological Monographs 94: e1635.

Petrén H, Köllner TG, Junker RR. 2023. Quantifying chemodiversity considering biochemical and structural properties of compounds with the R package chemodiv. New Phytologist 237: 2478–2492.

R Core Team. 2025. R: A Language and Environment for Statistical Computing. [WWW document] URL https://www.R-project.org/ [accessed 23 May 2026].

Ripley BD, Venables WN. 2002. Modern Applied Statistics with S, Fourth edition. New York: Springer.

Sasidharan R, Grond SG, Champion S, Eilers EJ, Müller C. 2024. Intraspecific plant chemodiversity at the individual and plot levels influences flower visitor groups with consequences for germination success. Functional Ecology 38: 2665–2678.

Scheriau CL, Nuerk NM, Sharbel TF, Koch MA. 2017. Cryptic gene pools in the *Hypericum perforatum*-*H. maculatum* complex: diploid persistence versus trapped polyploid melting. Annals of Botany 120: 955–966.

Schweiger R, Padilla-Arizmendi F, Nogueira-Lopez G, Rostás M, Lawry R, Brown C, Hampton J, Steyaert JM, Müller C, Mendoza-Mendoza A. 2021. Insights into metabolic changes caused by the *Trichoderma virens*–maize root interaction. Molecular Plant-Microbe Interactions 34: 524–537.

Shannon P, Markiel A, Ozier O, Baliga NS, Wang JT, Ramage D, Amin N, Schwikowski B, Ideker T. 2003. Cytoscape: a software environment for integrated models of biomolecular interaction networks. Genome Research 13: 2498–2504.

Silva BA, Ferreres F, Malva JO, Dias AC. 2005. Phytochemical and antioxidant characterization of *Hypericum perforatum* alcoholic extracts. Food Chemistry 90: 157–167.

Soltis PS, Liu X, Marchant DB, Visger CJ, Soltis DE. 2014. Polyploidy and novelty: Gottlieb’s legacy. Philosophical transactions of the Royal Society of London. Series B, Biological sciences 369: 20130351.

Speed MP, Fenton A, Jones MG, Ruxton GD, Brockhurst MA. 2015. Coevolution can explain defensive secondary metabolite diversity in plants. New Phytologist 208: 1251–1263.

Tavan M, Mirjalili MH, Karimzadeh G. 2015. In vitro polyploidy induction: changes in morphological, anatomical and phytochemical characteristics of *Thymus persicus* (Lamiaceae). *Plant Cell*, Tissue and Organ Culture 122: 573–583.

Thévenot EA, Roux A, Xu Y, Ezan E, Junot C. 2015. Analysis of the human adult urinary metabolome variations with age, body mass index and gender by implementing a comprehensive workflow for univariate and OPLS statistical analyses. Journal of Proteome Research 14: 3322–3335.

Thon FM, Müller C, Wittmann MJ. 2024. The evolution of chemodiversity in plants-From verbal to quantitative models. Ecology Letters 27: e14365.

Wetzel WC, Whitehead SR. 2020. The many dimensions of phytochemical diversity: linking theory to practice. Ecology Letters 23: 16–32.

Wickham H. 2016. ggplot2: Elegant Graphics for Data Analysis. Springer-Verlag New York.

Wickham H, Averick M, Bryan J, Chang W, McGowan L, François R, Grolemund G, Hayes A, Henry L, Hester J et al. 2019. Welcome to the Tidyverse. Journal of Open Source Software 4: 1686.

Wickham H, François R, Henry L, Müller K. 2026. dplyr: A Grammar of Data Manipulation. [WWW document] URL https://dplyr.tidyverse.org. [accessed 23 May 2026].

Wood TE, Takebayashi N, Barker MS, Mayrose I, Greenspoon PB, Rieseberg LH. 2009. The frequency of polyploid speciation in vascular plants. Proceedings of the National Academy of Sciences of the United States of America 106: 13875–13879.

Wu T, Bafort Q, Mortier F, Almeida-Silva F, Natran A, van de Peer Y. 2024. The immediate metabolomic effects of whole-genome duplication in the greater duckweed, *Spirodela polyrhiza*. American Journal of Botany 111: e16383.

Xiao X, Dussarrat T, Ziaja D, Seymen YB, Brokate L, Jakobs R, Weber B, Winkler JB, Schnitzler J-P, Müller C. 2025. Plastic Responses to Single and Combined Environmental Stresses in a Highly Chemodiverse Aromatic Plant Species. Physiologia Plantarum 177: e70626.

Xu Y, Jia H, Tan C, Wu X, Deng X, Xu Q. 2022. Apomixis: genetic basis and controlling genes. Horticulture Research 9: uhac150.

Yurekten O, Payne T, Tejera N, Amaladoss FX, Martin C, Williams M, O’Donovan C. 2024. MetaboLights: open data repository for metabolomics. Nucleic Acids Research 52: D640–D646.

