## Supplemental Figure 1 and 2 for "Evolutionary history and polyploidization lead to rapid shifts in chemodiversity of *Hypericum*"

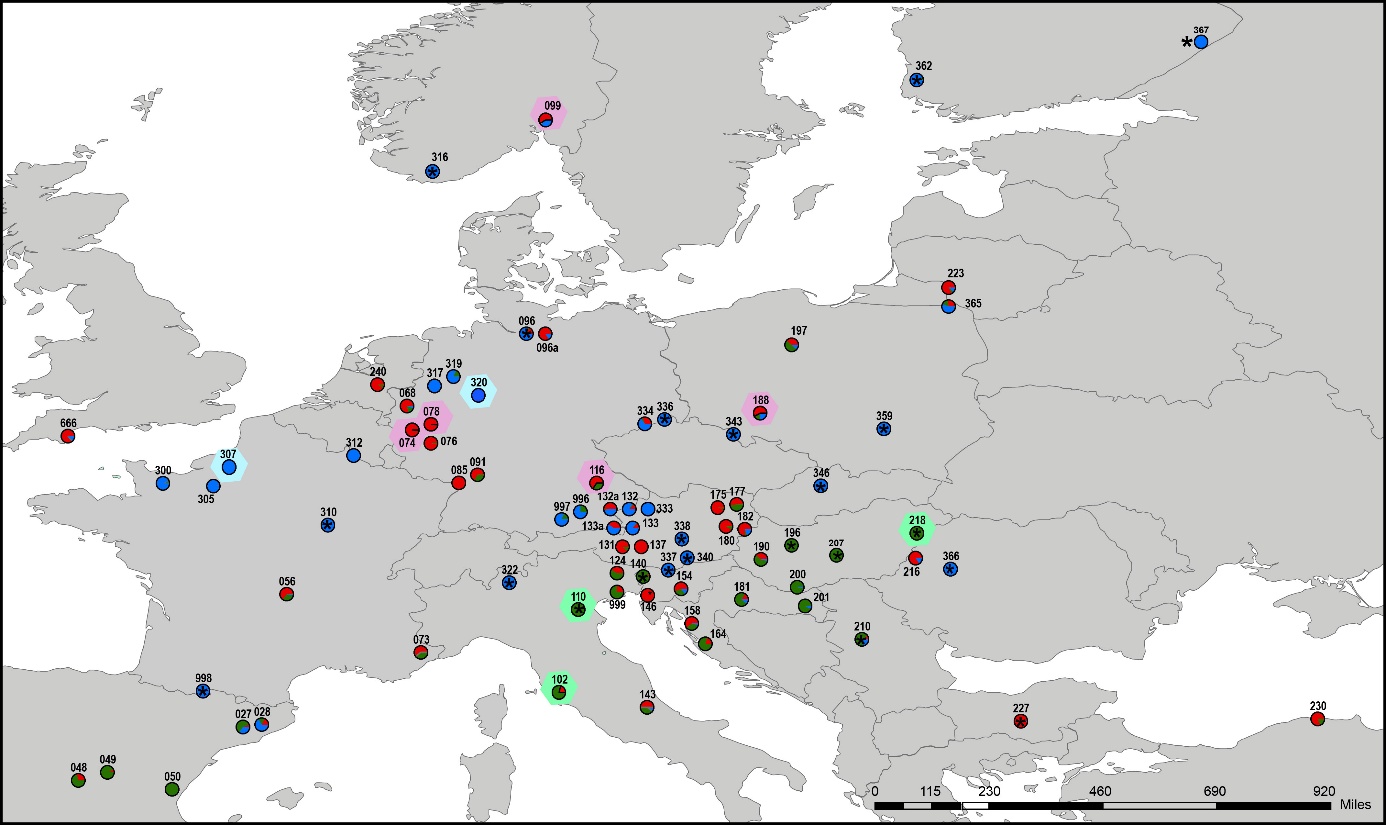
**Figure S1** Distribution map of geographically distinct *Hypericum* spp. populations, map modified from Koch *et al.* (2013). The population code is given above the pie charts which summarize population compositions. Asterisks (*) indicate diploid populations. blue = Hypericum maculatum; red = H. perforatum subsp. *perforatum*; green = H. perforatum subsp. *verononse*. Populations enclosed by hexagons were included in the present study.

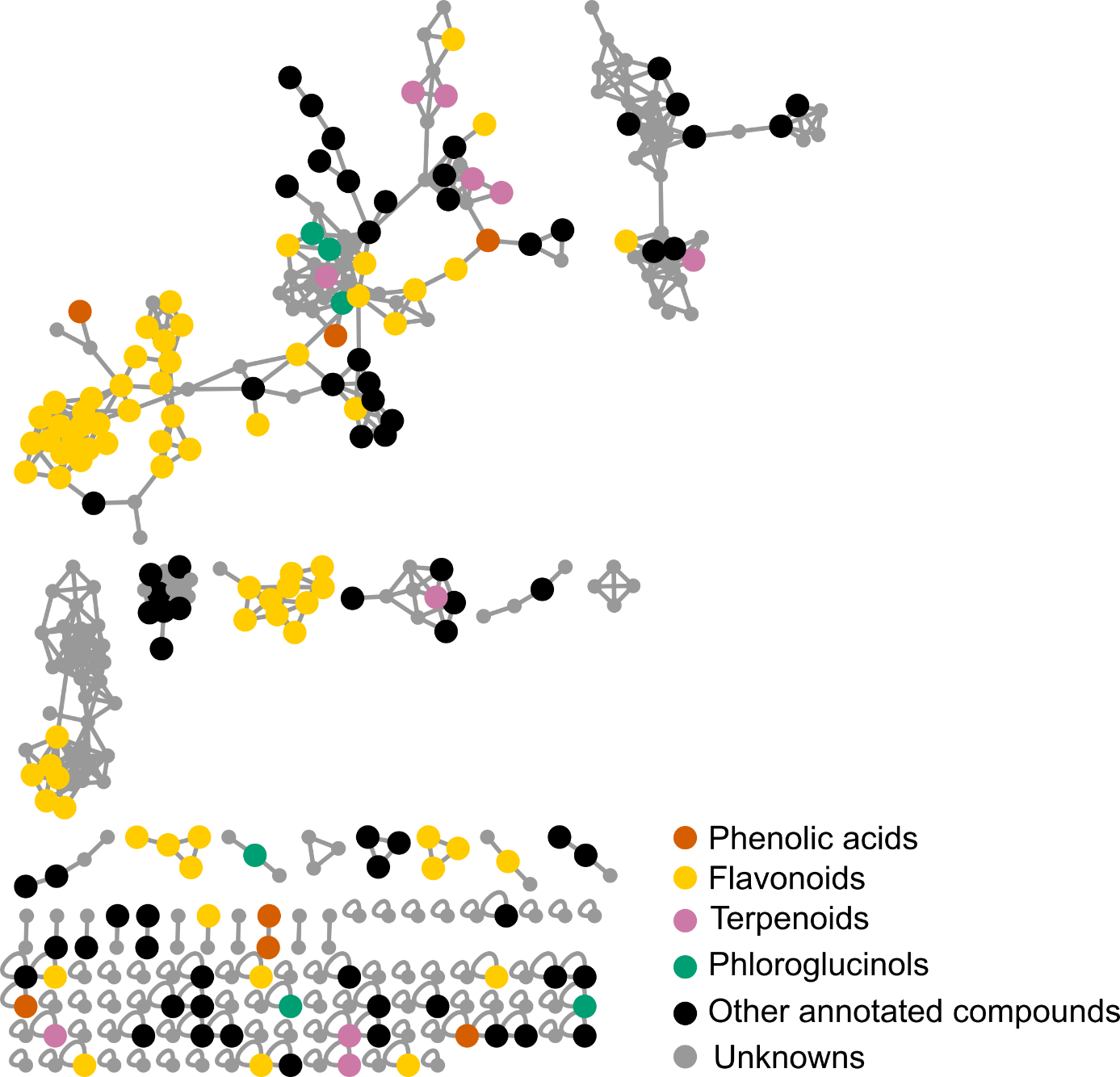

**Figure S2** Feature-based molecular network constructed from UHPLC-QToF-MS/MS electrospray ionization negative-mode data of leaf metabolic features in F1 offsprings of *Hypericum maculatum, H. perforatum* subsp. *perforatum*, *and H. perforatum* subsp*. veronense*. Features (nodes), identified using SIRIUS (with confidence score ≥ 0.8), were colored according to chemical families. Edges represent similarities in MS/MS fragmentation patterns, providing a visual measure of structural relatedness among features.

**References**

**Koch** **MA, Scheriau** **C, Betzin** **A, Hohmann** **N, Sharbel** **TF. 2013.** Evolution of cryptic gene pools in *Hypericum perforatum*: the influence of reproductive system and gene flow. *Annals of Botany* **111**: 1083-1094.
